# The Mediator tail module cooperates with proneural factors to determine fate identity in Glioblastoma

**DOI:** 10.1101/2021.05.04.442579

**Authors:** Moloud Ahmadi, Graham MacLeod, Nishani Rajakulendran, Milena Kosic, Andy Yang, Fatemeh Molaei, Sichun Lin, Zachary Steinhart, Nicole I. Park, Peter B. Dirks, Stephane Angers

## Abstract

Glioblastoma stem cells (GSCs) exhibit latent neuronal lineage differentiation potential governed by the proneural transcription factor Achaete-scute homolog 1 (ASCL1) and harnessing and promoting terminal neuronal differentiation has been proposed as a novel therapeutic strategy. Here, using a genome-wide CRISPR suppressor screen we identified genes required for ASCL1-mediated neuronal differentiation. This approach revealed a specialized function of the Mediator complex tail module and of its subunits MED24 and MED25 for this process in GSCs, human fetal neural stem cells and pluripotent stem cells. We show that upon induction of neuronal differentiation MED25 is recruited to genomic loci co-occupied by ASCL1 to regulate neurogenic gene expression programs. MED24 and MED25 are sufficient to induce neuronal differentiation in GSC cultures and to mediate neuronal differentiation in multiple contexts. Collectively our data expand our understanding of the mechanisms underlying directed terminal neuronal differentiation in brain tumor stem cells and point to a unique function of the Mediator tail in neuronal reprogramming.

## Introduction

One important barrier to Glioblastoma (GBM) treatment is the complex intratumoral heterogeneity of this cancer (Brennan et al., 2013; Neftel et al., 2019; Richards et al., 2021; Verhaak et al., 2010). Governing GBM growth are the self-renewing glioblastoma stem cells (GSC) that have tumor propagating properties with conserved, albeit defective, differentiation potential. GSCs are also resistant to chemoradiation and therefore underlie disease recurrence (Bao et al., 2006; Chen et al., 2012; Singh et al., 2004). GSCs phenotypically and functionally resemble neural precursor cells and are thought to hijack normal differentiation programs to gain self-renewal capacity (Chen et al., 2012). Recent single-cell RNA sequencing of tumor samples and cultured GSCs has revealed a transcriptional gradient in GSCs between states characterized by neurodevelopment and inflammatory/injury-response processes (Richards et al., 2021; Wang et al., 2019b). The functional resemblance of tumor cells to normal brain stem/precursor cells has led to the hypothesis that inducing differentiation into non-proliferative central nervous cell types could represent a therapeutic strategy for GBM (Carén et al., 2015; Park et al., 2017; Piccirillo et al., 2006). GSCs induced to differentiate into astrocytes and oligodendrocyte-like cells using BMP have been shown to be vulnerable to cell-cycle re-entry (Carén et al., 2015). In contrast, neurons represent a terminally differentiated cell type that studies have shown are resistant to transformation (Alcantara Llaguno et al., 2019). Thus, manipulation of the epigenetic and transcriptional landscapes of GSCs to promote their terminal differentiation into a committed neuronal state, is predicted to reduce tumor proliferative capacity and increase survival.

The acquisition of neuronal identity is driven by the coordinated activity of various proneural transcription factors (TFs), activators and repressors that integrate spatial and temporal cues from the environment. The TFs belonging to the proneural basic Helix-Loop-Helix (bHLH) family are cell fate instructors, expressed during embryonic development and throughout life in regions of the brain where neural stem cell (NSC) niches exist (Bertrand et al., 2002; Shirasaki and Pfaff, 2002). These proneural proteins participate in neuronal commitment, cell cycle exit and ultimately neuron formation. For instance, induced expression of the neurogenic bHLH transcription factor Achaete-scute homolog 1 (ASCL1) has been shown to promote neuronal differentiation in multiple cell types, functioning as a pioneer factor opening closed regions on chromatin and thereby activating a neurogenic transcriptional program (Chanda et al., 2014; Raposo et al., 2015; Wapinski et al., 2017). Across GSCs, high *ASCL1* expression defines a subset with latent capacity for terminal neuronal differentiation (Park et al., 2017). Inhibition of Notch and/or Wnt signaling in these cells or forced overexpression of *ASCL1* invoke robust neuronal differentiation and reduce clonogenic capacity (Park et al., 2017; Rajakulendran et al., 2019). However, how neurogenic factors evoke cell cycle exit in stem-like cells and drive commitment to a neuronal state is not well characterized.

An additional level of regulation of gene expression is achieved by transcriptional co-regulators. The Mediator is an evolutionarily conserved multisubunit protein complex that functions as a transcriptional co-activator and serves as a physical and functional interface between DNA-bound TFs and RNA polymerase II (Pol II) (Allen and Taatjes, 2015; Yin and Wang, 2014). Mediator comprises four modules: head, middle, tail and a kinase domain (Conaway and Conaway, 2013). The head module binds the Pol II apparatus and the tail module interacts with transcription factors. Collectively, the head, middle and tail modules form what is known as the Mediator core complex (Sierecki, 2020). The kinase module has a regulatory function and binds to the core in a reversible manner (Andrau et al., 2006; Knuesel et al., 2009). Mediator not only acts in transcriptional machinery assembly and stabilization, but it has also been described as important for DNA bending, chromatin remodeling, and nuclear gene positioning (Fournier et al., 2016; Lai et al., 2013; Soutourina, 2018). In the last few years, Mediator has been localized at super-enhancers and hence linked to control of cell fate identity and tumorigenesis (Schiano et al., 2014; Whyte et al., 2013; Yin and Wang, 2014).

In this study, using an unbiased genome*-*wide CRISPR*-*Cas9 suppressor screen we determined that members of the Mediator complex belonging to the tail module while dispensable for GSC growth, are required for their neural commitment and differentiation. Particularly, through integration of gene expression data with chromatin immunoprecipitation sequencing (ChIP-seq), we found that ASCL1 and Mediator tail subunit MED25 mutually interact with specific chromatin regions to direct changes in transcriptional programs that culminate in the adoption of neuronal cell identity. Our findings, although uncovered in the context of GSCs, also point to a role for the Mediator tail domain for neuronal differentiation in human embryonic stem cells suggesting a broader role during neuronal reprogramming and potentially underlying its deficiencies in neurodevelopmental disorders (Bernard et al., 2006; Hashimoto et al., 2011; Leal et al., 2009).

## Results

### MED24 and MED25 are required for ASCL1-mediated neuronal differentiation

We previously established that *ASCL1* overexpression is sufficient to trigger neuronal differentiation in Glioblastoma stem cells and hence leads to proliferation arrest, decreased self-renewal and inhibition of tumorigenicity (Park et al., 2017). To identify genes required for ASCL1-mediated neuronal differentiation in this context, we first introduced a doxycycline (DOX)-inducible *ASCL1* cassette into the G523 patient-derived developmental-subtype GSC culture: G523-Tet-ON-ASCL1 (Figure 1A). Treatment of these cells with DOX for two weeks led to *ASCL1* overexpression and resulted in neuronal lineage differentiation, determined by a significant increase in ßIII-tubulin (TUBBIII) positive cells, and as a result led to a decrease in proliferation marker Ki67 (Figures 1B).

**Figure 1.**
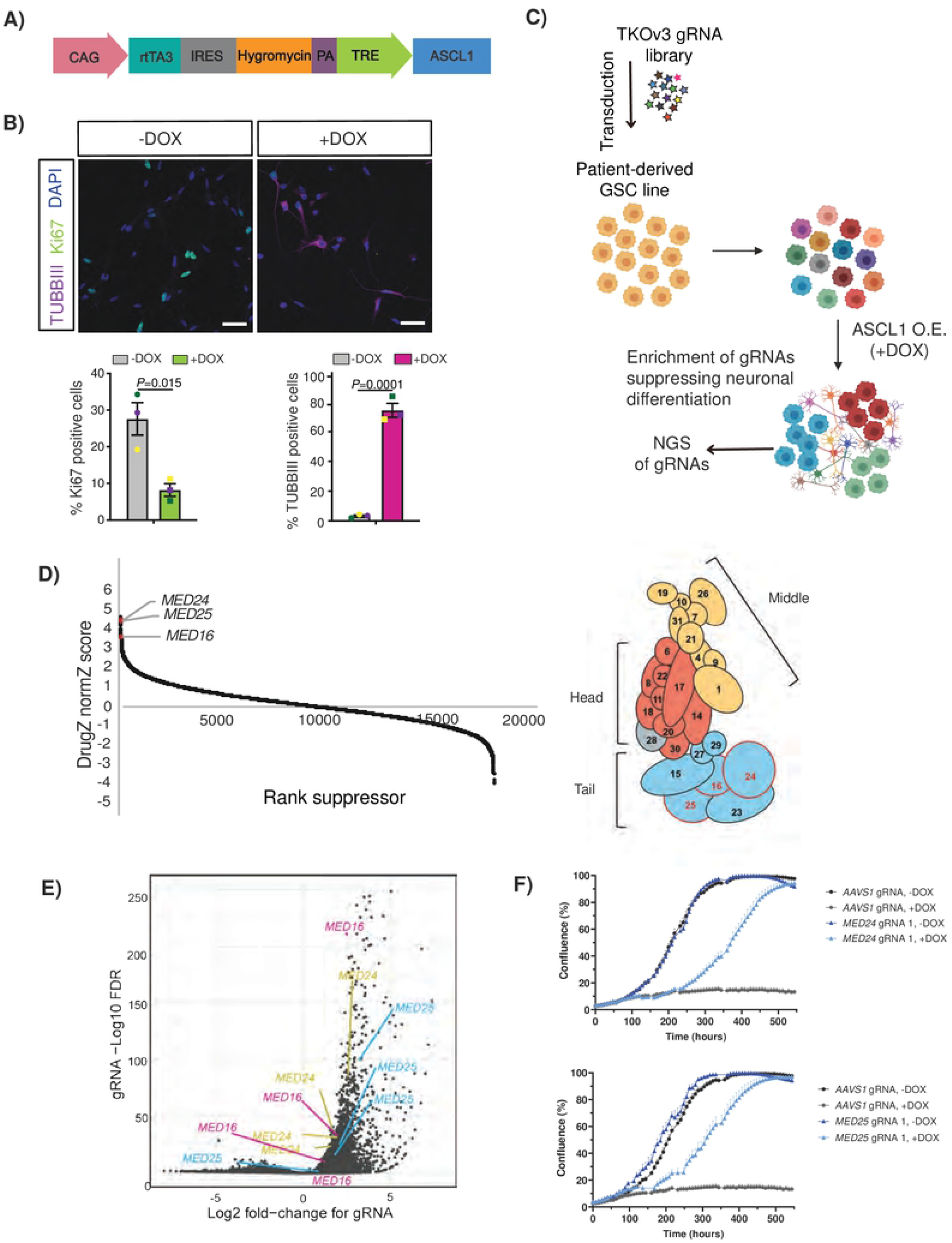
Genome-wide CRISPR-Cas9 screen identifies novel hits required for ASCL1-mediated effect on cell fate commitment. (A) Tet-ON-ASCL1 construct used to generate the G523-Tet-ON-ASCL1 cell line. (B) Immunocytochemical staining (top) and quantification (bottom) of TUBIII and Ki67 positive cells in G523-Tet-ON-ASCL1 GSCs treated with DOX for 2 weeks. Cell nuclei were stained with DAPI. Scale bar, 50 μm. Data represents mean ± SEM (n=3, color-coded). Statistics are derived from a two-tailed unpaired t test. Error bars, mean ± SEM. (C) Schematic of genome-wide CRISPR screens for identification of ASCL1-mediated targets driving neuronal differentiation in GSCs. (D) Left: Rank order plot of DugZ-calculated normZ scores from genome-wide CRISPR screen in G523-Tet-ON-ASCL1 GSCs. Ranks based on suppression of growth arrest. Identified MED subunits are highlighted. Right: Diagram of the mammalian mediator complex depicting the head (red), middle (yellow) and tail (modules) and subunits identified in CRISPR-Cas9 screen (red text). MED28 has not been assigned to a specific module. Modified from (Soutourina, 2018). (E) Volcano plot of gRNA-level enrichment p-values from genome-wide CRISPR screen calculated by the MAGeCK algorithm. Identified MED subunits are highlighted. (F) Quantification of cell confluency of AAVS1KO, MED24KO (top) and MED25KO (bottom) G523-Tet-ON-ASCL1 GSCs in the presence or absence of DOX (n=3). Error bars, mean ± SD. See also Figure S1 and Supplemental File-1

Since DOX-induced *ASCL1* overexpression leads to proliferation arrest, we predicted that knockout of genes required for this neuronal induction circuit would confer a proliferative advantage over differentiating cells. With this in mind, we performed a genome-wide CRISPR-Cas9 suppressor screen using the TKOv3 gRNA library in the presence of DOX for four weeks (Figure 1C) (Hart et al., 2017; MacLeod et al., 2019). Using the DrugZ algorithm (Colic et al., 2019) we identified 30 genes (FDR < 0.1) whose knockout suppresses ASCL1-mediated proliferation arrest (Figure 1D and Supplemental Data file-1). Intriguingly, 3 of our hits (*MED16, MED24* and *MED25)* encode tail module subunits of the Mediator Complex (Figure 1D). A second algorithm, MAGeCK (Li et al., 2014) also identified *MED24* and *MED25* within the top 5 hits in the screen while *MED16* was the 15th most positively selected gene (Figure 1E, Supplemental Data file-1). Therefore, we validated the requirement for MED24 and MED25 in ASCL1-dependent arrest of GSC proliferation by transducing G523-Tet-ON-ASCL1 cells with lentivirus expressing individual targeting gRNAs (Figure S1A). Cells expressing control gRNA (AAVS1KO) or gRNA targeting *MED24 (*MED24KO) or *MED25* (MED25KO) were then treated with DOX for 2 weeks to initiate ASCL1-mediated neuronal differentiation. While ASCL1 over-expression resulted in proliferation arrest in control cells, targeted perturbation of *MED24* or *MED25* using two individual gRNAs robustly abrogated this effect (Figures 1F and Figure S1B-C). Given the importance of the Mediator Complex in regulation of RNA Pol II-mediated gene transcription (El Khattabi et al., 2019; Levine et al., 2014), we examined if suppression of the ASCL1-mediated decrease in growth, observed in *MED24* and *MED25* knockout cells, was simply due to inhibition of the inducible expression cassette. Therefore, we assessed *ASCL1* mRNA levels in control, MED24KO and MED25KO GSCs line upon 14 days of DOX treatment. This experiment revealed similar *ASCL1* expression in both MED24KO and MED25KO cells suggesting that these Mediator subunits are dispensable for *ASCL1* expression (Figure S1D) and is consistent with the tail module not being required for the core transcriptional activity of Mediator.

Given that the readout of our CRISPR-Cas9 screen was proliferation and not a direct assessment of differentiation, we next tested if *MED24* or *MED25* deletion leads to defective ASCL1-mediated neuronal differentiation. Following *ASCL1* induction, Immunofluorescence imaging of MED24KO and MED25KO cells revealed a decrease in TUBBIII positivity with a concomitant increase in Ki67 positivity when compared to control cells (Figure 2A), indicating a diminished capacity for differentiation towards the neuronal lineage in the absence of these Mediator subunits. Neuronal identity is associated with decreased expression of stem cell markers such as SOX2 and Nestin (Graham et al., 2003; Park et al., 2017). Given the decreased ASCL1-dependent neuronal differentiation of G523 GSCs harboring *MED24* or *MED25* deletion, we asked if these cells retained expression of Nestin following ASCL1 induction. Consistent with the requirement of *MED24* and *MED25* in ASCL1-mediated neuronal differentiation, we observed that while control cells showed a significant decrease in Nestin expression following ASCL1 induction, MED24KO and MED25KO cells displayed Nestin levels similar to uninduced GSCs (Figures 2B and 2C). We next asked whether the requirement for MED24 and MED25 in ASCL1-mediated neuronal differentiation also extended more broadly to gliogenesis potential by assessing astrocytic differentiation. We observed no change in the expression of the astrocytic marker glial fibrillary acidic protein (GFAP) either at the basal level or under ASCL1 over-expression indicating that loss of neuronal differentiation upon MED24 or MED25 depletion was not due to concomitant increases in astrocytic differentiation (Figure 2D). We were also interested to determine whether depletion of these mediator subunit genes also affected self-renewal capacity. Interestingly, we found that while ASCL1 over-expression markedly blocked self-renewal capability of GSCs, with only 0.3 % of cells capable of initiating a colony, MED24KO and MED25KO abrogated this effect resulting in higher sphere forming potential (Figure 2E). Collectively, these experiments establish a required role for Mediator Complex subunits MED24 and MED25 in ASCL1-mediated neuronal differentiation of GSCs.

**Figure 2.**
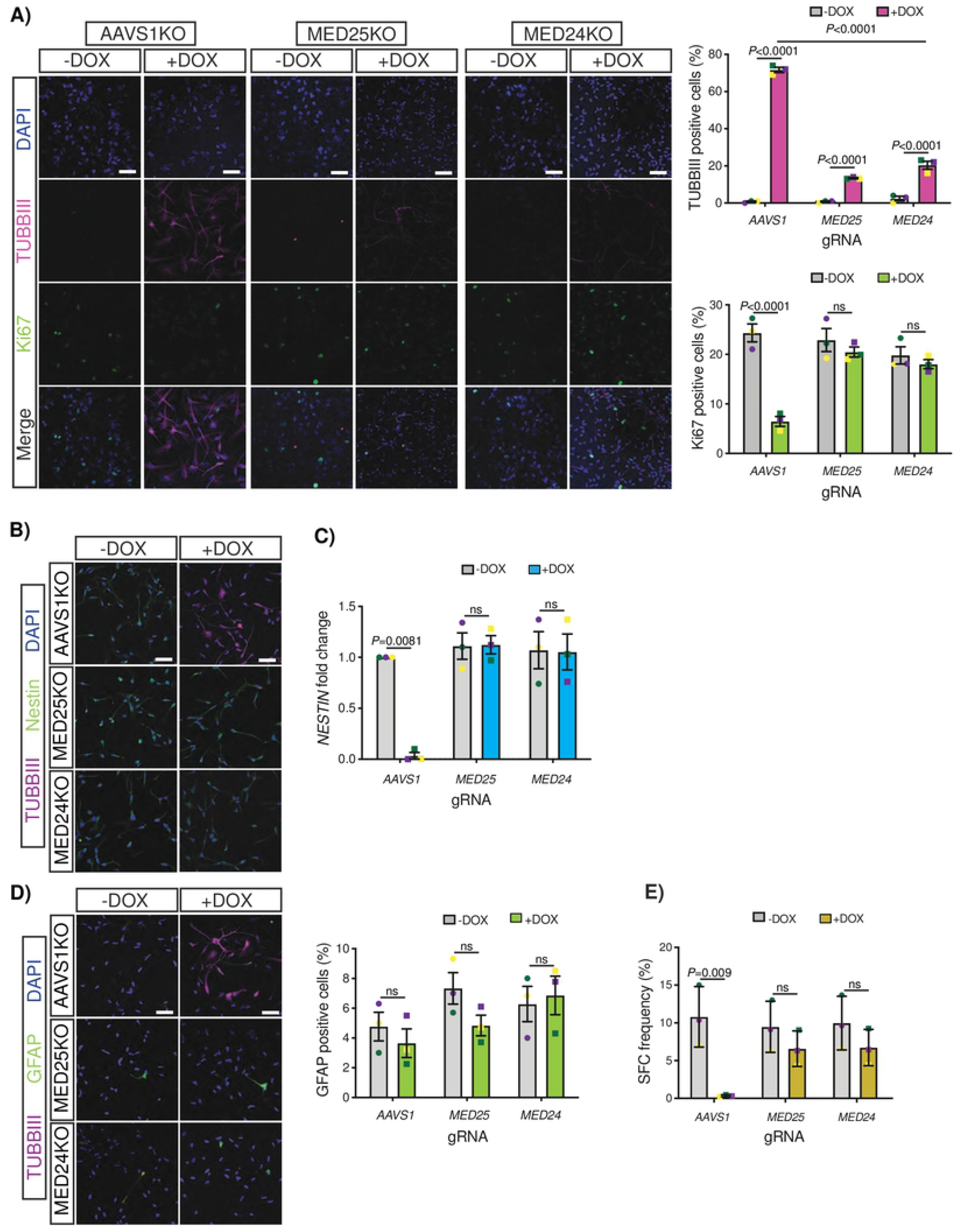
MED24-MED25 complex is required for ASCL1-mediated neuronal differentiation. (A) Representative immunocytochemical staining (left) and quantification (right) of TUBBIII and Ki67 of AAVS1KO, MED24KO and MED25KO G523-Tet-ON-ASCL1 GSCs induced with DOX for 2 weeks. Cell nuclei were stained by DAPI. Scale bar, 50 μm. Shown are mean values ± SEM (n=3, color-coded) tested by two-way ANOVA and Tukey’s multiple comparison tests. (B) Representative immunocytochemical staining of TUBBIII and Nestin positive cells of AAVS1KO, MED24KO and MED25KO G523-Tet-ON-ASCL1 GSCs induced with DOX for 2 weeks. Cell nuclei were stained by DAPI. Scale bar, 50 μm. (C) qPCR analysis of Nestin mRNA levels of AAVS1KO, MED24KO and MED25KO G523-Tet-ON-ASCL1 GSCs treated with or without DOX (n=3, color-coded). two-way ANOVA and Bonferroni’s multiple comparison tests. Error bars, mean ± SD. (D) Representative immunocytochemical staining (left) and quantification (right) of TUBBIII and GFAP positive cells of AAVS1KO, MED24KO and MED25KO G523-Tet-ON-ASCL1 GSCs induced with DOX for 2 weeks. Cell nuclei were stained by DAPI. Scale bar, 50 μm. Shown are mean values ± SEM (n=3, color-coded) tested by two-way ANOVA and Tukey’s multiple comparison tests. (E) *In vitro* LDA of AAVS1KO, MED24KO and MED25KO G523-Tet-ON-ASCL1 GSCs treated with or without DOX. Error bars, estimated frequency ±95% CI (n=5, color-coded). two-way ANOVA and Tukey’s multiple comparison tests.

### MED24 and MED25 govern gene expression required for ASCL1-mediated neuronal differentiation

The Mediator’s role as regulator of transcription prompted us to next examine the downstream targets of MED25 that mediate neuronal differentiation in response to ASCL1 over-expression (Figures 3A-3B). We first performed RNA sequencing (RNA-seq) analysis of G523-Tet-ON-ASCL1 GSCs following seven days of *ASCL1* induction, identifying 1,937 differentially expressed genes (Figure 3A). Conversely, in MED25KO cells there were proportionately less transcriptional changes following ASCL1 induction (390 genes; Figure 3B) suggesting suppression of the neuronal induction response (Figure 3C). A common set of 187 genes were differentially expressed following DOX treatment in both control and MED25KO cells indicating the existence of ASCL1 induced transcriptional programs independent of MED25. Strikingly, genes differentially expressed between control and MED25KO cells following ASCL1 induction are enriched for functions in neurogenesis and nervous system development explaining abrogation of ASCL1-mediated neuronal conversion in the absence of MED25 (Figure 3D). A more detailed examination of these genes revealed multiple neuronal genes in which transcriptional response to ASCL1 overexpression was abolished in the absence of MED25 (Figure 3E and Supplemental Figure 2). In addition, we observed that genes from multiple signaling pathways involved in self-renewal and proliferation, such as Wnt, Notch, Shh and FGF signaling, were upregulated upon MED25 loss (Figure 3E and Supplemental Figure 2). This data supports specific roles for MED25 in the suppression of self-renewal and promotion of neuronal differentiation. qRT-PCR confirmed that previously known ASCL1 targets (Park et al., 2017) *DLL3* and *TMPRSS2* were downregulated upon *MED24* and *MED25* knockout further suggesting the importance of these Mediator Complex tail subunits in modulation of ASCL1-mediated neurogenic programs (Figure 3F).

**Figure 3.**
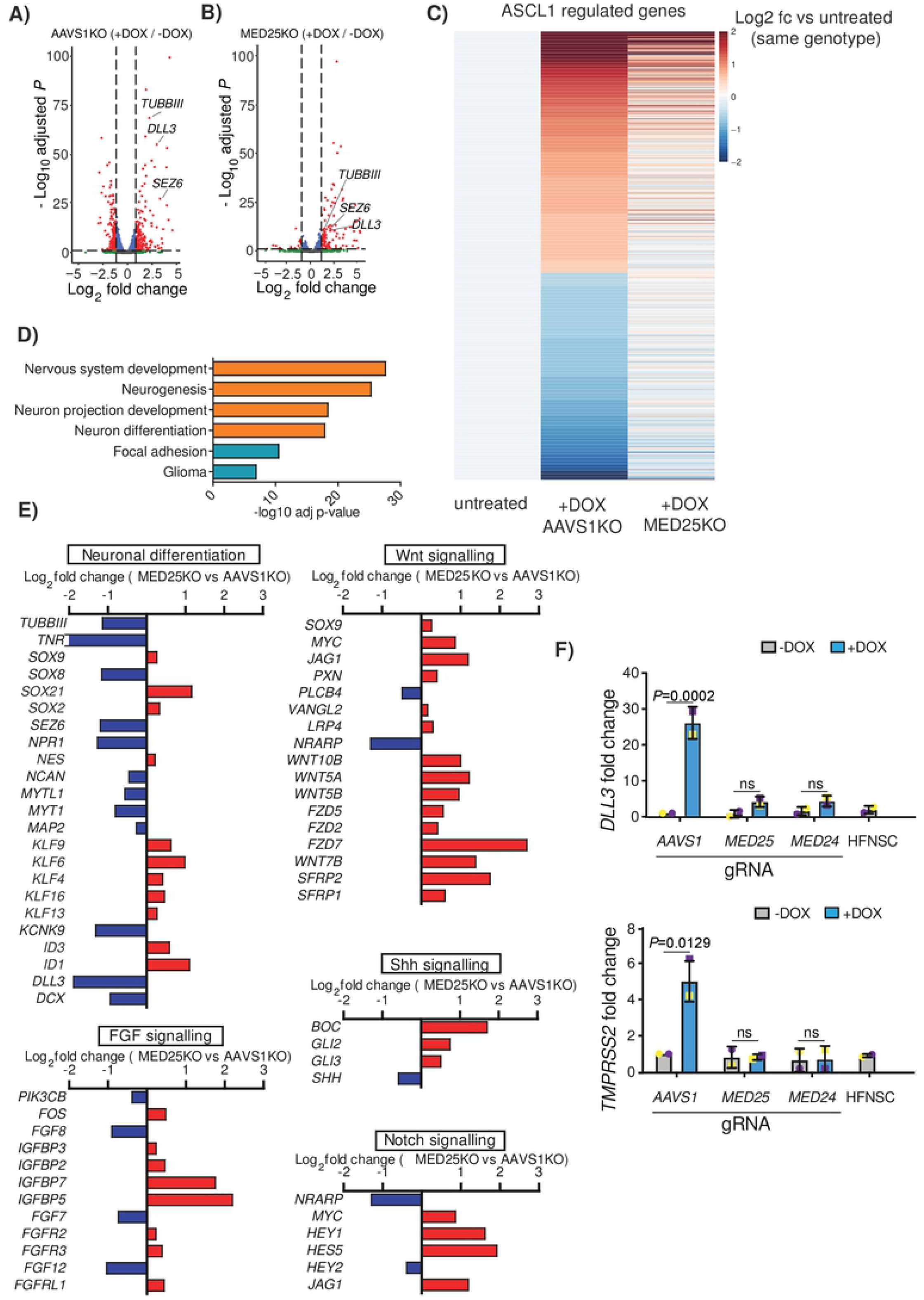
RNA-Seq analysis uncovers MED25 neuronal target genes upon ASCL1 over-expression. (A) Volcano plot depicting differentially expressed genes from RNA-seq in AAVS1KO (+DOX) relative to AAVS1KO(-DOX) G523-Tet-ON-ASCL1 GSCs induced with DOX for 1 week (n=3). (B) Volcano plot depicting differentially expressed genes from RNA-seq in MED25KO (+DOX) relative to MED25KO(-DOX) G523-Tet-ON-ASCL1 GSCs induced with DOX for 1 week (n=3). (C) Heatmap of average transcriptional response (n=3) to DOX-induced ASCL1 overexpression in AAVS1KO and MED25KO relative to their respective uninduced control samples. Color scale = log2 fold-change from -DOX. (D) Selected Gene ontology biological processes (orange) and KEGG pathways (cyan) enriched in differentially expressed genes between AAVS1KO control and MED25KO G523-Tet-ON-ASCL1 GSCs induced with DOX for 1 week. (E) Gene expression changes in MED25KO (+DOX) vs. AAVS1KO (+DOX) control cells in selected signalling pathways from RNA-seq experiments. (F) qPCR analysis of mRNA levels of differentially expressed genes in AAVS1KO, MED24KO and MED25KO G523-Tet-ON-ASCL1 GSCs treated with or without DOX and normalized to AAVS1KO (-DOX) (n=2, color-coded). two-way ANOVA and Bonferroni’s multiple comparison tests. Error bars, mean ± SD. See also Figure S2 and Supplemental File-2

### MED25 directly binds to enhancer regions corresponding to neuronal target genes

To determine whether MED25 directly occupies regulatory regions controlling the neuronal genes identified in our RNA-seq analysis, we performed MED25 chromatin immunoprecipitation sequencing (ChIP-seq) of G523-Tet-ON-ASCL1 GSCs after 18 hours of DOX induction triggering ASCL1-mediated differentiation. MED25KO cells served as a negative control for ChIP using the MED25 antibody. We identified 3,785 regions bound by MED25 that consisted primarily of sites distal to the transcription start site (TSS). Only 2% and 8% of bound regions correspond to exonic and promoter regions respectively (Figures 4A and 4B, Supplemental data file 3). Next, the distal MED25 bound regions were compared to databases of annotated cis-regulatory elements (CREs). We found that 15% of MED25 bound regions overlapped with known enhancers (3% known to be active in neural stem and precursor cells) and 4% overlapped with super-enhancers identified in brain-derived cell lines (Gao and Qian, 2020; Qian et al., 2019; Wang et al., 2019a) (Figure 4B). Analysis of genes proximal to MED25-bound regions revealed enrichment for GO Biological Processes associated with neurogenesis and nervous system development further pointing to the engagement of MED25 in ASCL1-mediated neuronal reprogramming (Figure 4C and 4D). These neuronal genes included *TUBB3, NEUROG3, MTY1L* and *KCNK9* that were also found to be induced by ASCL1 overexpression (Figure 3) and (Park et al., 2017). Motif enrichment analysis of MED25-bound regions showed a modest but significant (q-value = 0.0038) enrichment for the E-box sequence CACGTG, which has been identified as a binding site for proneural transcription factors associated with mediator complex subunits in mouse neural stem cells (Quevedo et al., 2019) (Figure 4E).

**Figure 4.**
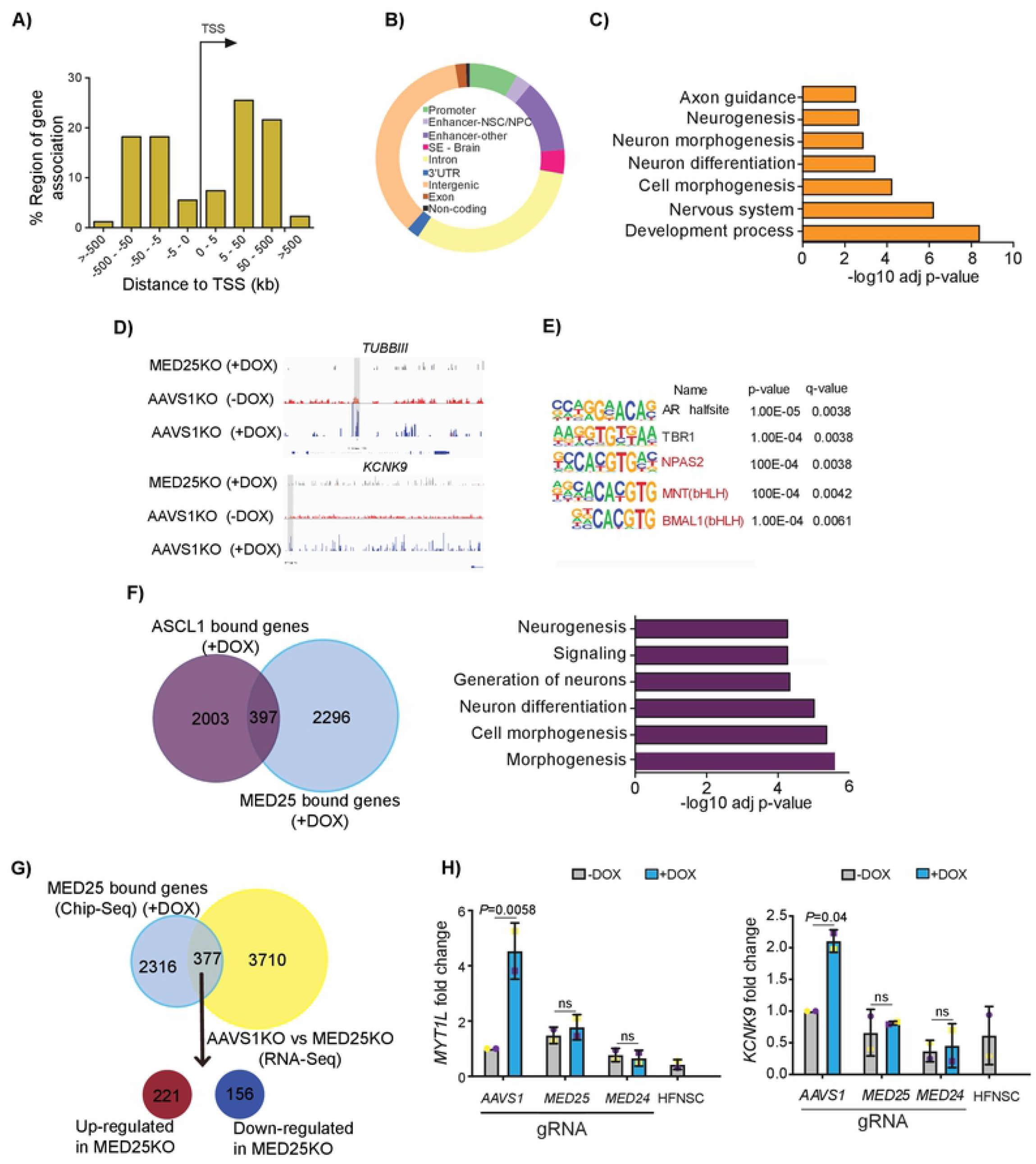
MED25 binds to various neuronal target genes upon ASCL1 over-expression. (A) Location of MED25 binding sites in relation to the closest annotated TSS (B) Doughnut displaying genomic distribution of MED25-binding sites (C) gProfiler analysis reveals that MED25 binds to genomic loci involved in neuronal differentiation and nervous system development. (D) Representative tracks of Med25 ChIP-seq peaks of AAVS1KO and MED25KO G523-Tet-ON-ASCL1 GSCs induced with DOX for 18 hours near the KCNK9 and TUBB3 loci (n=3). (E) Motif analysis for MED25 bound regions (F) Left: Venn diagram revealing the overlap between ASCL1 bound genes and MED25 bound genes in G523-Tet-ON-ASCL1 GSCs induced with DOX for 18 hours. Right: gProfiler analysis of the genes that are mutually bound by ASCL1 and MED25 upon ASCL1-induced neuronal differentiation. (G) Venn diagram revealing the overlap between MED25 bound gene loci and genes differentially upon ASCL1-induced neuronal induction in AAVS1KO versus MED25KO G523-Tet-ON-ASCL1 GSCs. (H) qPCR analysis of mRNA levels of differentially expressed genes in AAVS1KO, MED24KO and MED25KO G523-Tet-ON-ASCL1 GSCs treated with or without DOX and normalized to AAVS1KO (-DOX) (n=2, color-coded). two-way ANOVA and Bonferroni’s multiple comparison tests. Error bars, mean ± SD. See also Supplemental File-3.

We then compared MED25 occupied regions with previously reported ASCL1 bound regions in the same cellular model (Park et al., 2017). We found that 397 genes associated with MED25-bound regions were also occupied by ASCL1 (Figure 4F) and that these genes were associated with biological processes linked to nervous system development (Figure 4G). To gain additional insights into the role of MED25 in modulation of neural target gene expression, we analyzed the overlap of our RNA-seq data with our MED25 ChIP-Seq data and found 377 genes differentially expressed upon *MED25* deletion that were associated with MED25-bound regions (Figure 4H). This suggests that MED25 may directly modulate the identified genes as a consequence of ASCL1 over-expression. To verify that MED24 plays a similar role to MED25, as indicated by our CRISPR-screen, we tested the effect of individual mediator subunit depletion on the expression of *KCNK9* and *MYT1L*. Both were identified as differentially expressed in our RNA-seq experiment, and associated with MED25 bound regions according to our ChIPseq analysis. We found that deletion of either *MED24* or *MED25* led to an abrogation of ASCL1 stimulated increases in expression levels (Figures 4I). We conclude that MED24/25 tail subunits of Mediator are required for ASCL1-mediated gene expression regulation and neuronal differentiation.

### MED24 and MED25 are required for neuronal conversion of HFNS and ESCs

Thus far our results suggest an important role for MED24 and MED25 in governing the neuronal differentiation of GSCs upon ASCL1 over-expression by binding CREs and underlying the expression of ASCL1 neuronal target genes. Growth factor (GF) withdrawal has been shown to induce expression of neurogenic factors leading to neuronal conversion of GSCs and Human Fetal Neural Stem cells (HFNS) (Pollard et al., 2009). We therefore sought to determine whether MED24 and MED25 are required for neuronal differentiation outside the context of ASCL1 over-expression in GSCs. Consistent with our findings above, while control cells displayed increased staining for neuronal marker TUBB3 after GF withdrawal, MED24KO and MED25KO led to failure in neuronal differentiation in two different patient-derived GSC cultures (Figures 5A and S3A). Similarly, we observed that MED24 and MED25 also mediating neuronal differentiation of HFNS lines in response to growth factor removal (Figures 5B and S3B-D). These results suggest a broader role for MED24 and MED25 in neuronal differentiation outside the context of ASCL1 over-expression.

**Figure 5.**
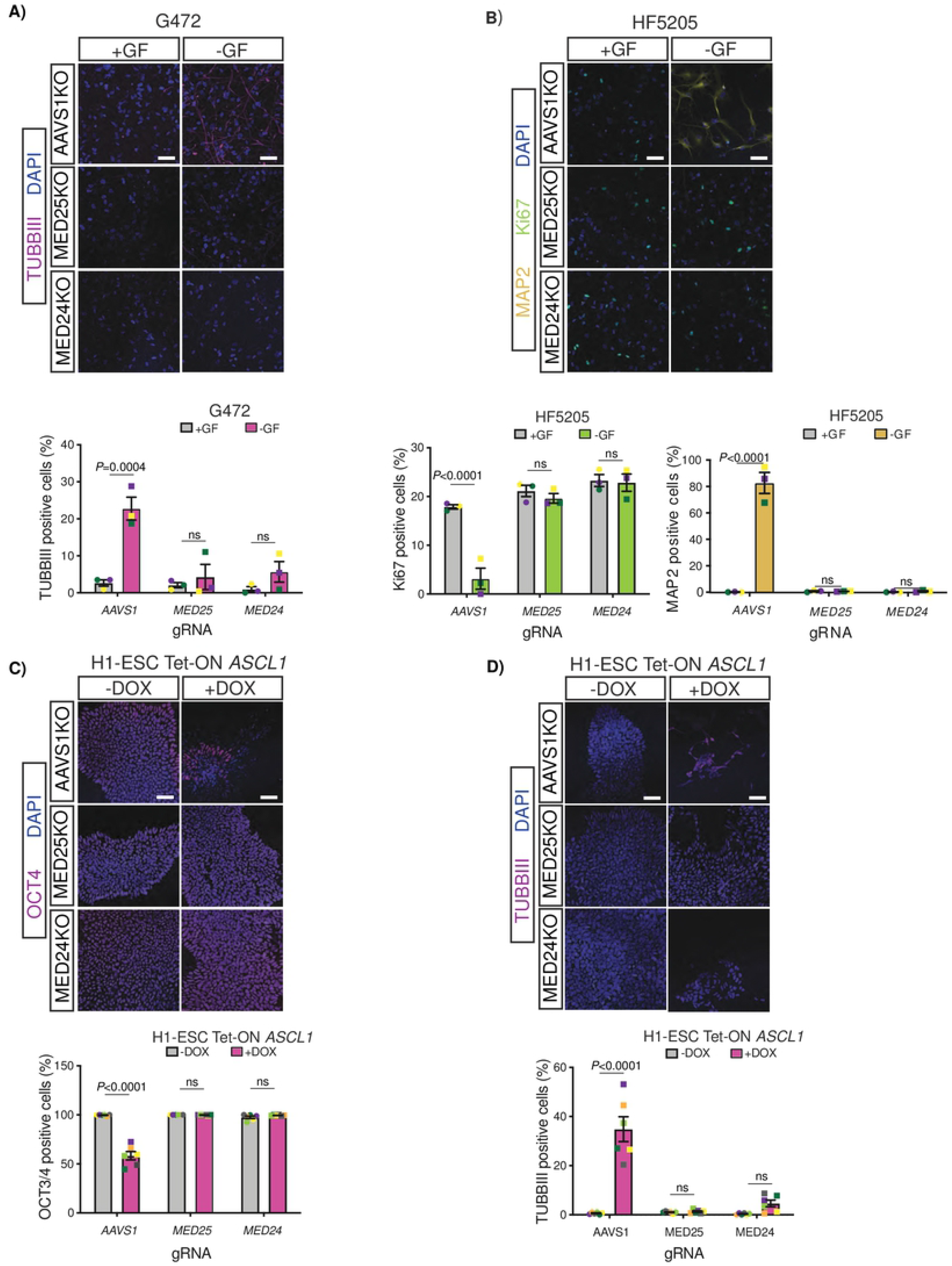
MED24 and MED25 are required for neuronal differentiation in HFNS, GSCs as well as ESCs. (A) Immunocytochemical staining (top) and quantification (bottom) of TUBBIII positive cells of AAVS1KO, MED24KO and MED25KO GSCs (G472) grown in the presence or absence of growth factor (GF). Cell nuclei were stained by DAPI. Scale bar, 50 μm. Shown are mean values ± SEM (n=3, color-coded) tested by two-way ANOVA and Tukey’s multiple comparison tests. (B) Immunocytochemical staining (top) and quantification (bottom) of Ki67 and MAP2 positive cells of AAVS1KO, MED24KO and MED25KO HFNS (HF5205) grown in the presence or absence of GF. Cell nuclei were stained by DAPI. Scale bar, 50 μm. Shown are mean values ± SEM (n=3, color-coded) tested by two-way ANOVA and Tukey’s multiple comparison tests. (C) Immunocytochemical staining (top) and quantification (bottom) of OCT4 positive cells of AAVS1KO, MED24KO and MED25KO ESCs-Tet-ON-ASCL1 cells induced with DOX for 6 days. Cell nuclei were stained by DAPI. Scale bar, 50 μm. Shown are mean values± SEM (n=3, color-coded) tested by two-way ANOVA and Tukey’s multiple comparison tests. (D) Immunocytochemical staining (top) and quantification (bottom) of TUBBIII positive cells of AAVS1KO, MED24KO and MED25KO ESCs-Tet-ON-ASCL1 cells induced with DOX for 6 days. Cell nuclei were stained by DAPI. Scale bar, 50 μm. Shown are mean values ± SEM (n=3, color-coded) tested by two-way ANOVA and Tukey’s multiple comparison tests. See also Figure S3.

ASCL1 over-expression was also shown to induce neuronal differentiation in ESCs (Chanda et al., 2014). Therefore, we next inquired whether MED24 and MED25 are required for neuronal fate commitment in ESCs in response to forced ASCL1 expression by introducing the inducible Tet-ON-ASCL1 construct (Figure 1A) into the H1 ESC line. After 6 days of DOX treatment, we observed a decrease in expression of the stemness marker OCT3/4 (Figures 5C and S3E) that was accompanied with an increase in expression of the neuronal marker TUBBIII in control cells (Figures 5D). However, this effect was abrogated in MED24KO and MED25KO ESCs suggesting the requirement of MED24 and MED25 in the ASCL1-mediated neuronal conversion in the context of ESCs. This data also indicates that MED24 and MED25 are dispensable for pluripotency maintenance in ESCs as their depletion did not influence OCT3/4 expression (Figures 5C and 5D).

### MED24 and MED25 are sufficient for neuronal differentiation and are required for neuronal conversion mediated by various neurogenic transcription factors

Given the requirement of MED24 and MED25 in neuronal conversion, we next sought to determine if their individual overexpression is sufficient to induce this process. We therefore, introduced the previously described dCAS9-VPR construct (Chavez et al., 2015) (Figure S4A) into the G523 GSC culture (GSC-dCAS9-VPR) to allow for targeted activation of gene expression. GSC-dCAS9-VPR cells were transduced with gRNAs individually targeting TSS proximal regions of the *GAL4* (negative control), *ASCL1* (positive control), *MED24* and *MED25* genes. Target gene activation was then induced for 14 days by DOX treatment. Strikingly, we observed that induction of *MED24* and *MED25* resulted in both a decrease in proliferation and a significant increase in the number of TUBBIII positive cells that displayed elongated, and extended neurites similar to mature neurons (Figures 6A-C). We confirmed via qRT-PCR, that robust increases in mRNA expression of *ASCL1, MED24* and *MED25* were evident following 14 days of dCAS9-VPR activation (Figure 6D). The degree of neuronal induction observed with *MED24* and *MED25* overexpression was less than that of *ASCL1;* nonetheless, it demonstrated a sufficiency of these Mediator subunits in this process (Figures 6A-C). Finally, we sought to gain a better understanding of whether the tail Mediator subunits also cooperate with other neurogenic transcription factors. We therefore generated MED24KO and MED25KO GSC-dCas9-VPR lines and transduced these cells with gRNAs individually targeting 1000-2000bp upstream of the TSS of *NEUROD1, NEUROD6* and *NGN1-NGN2* genes as well as *ASCL1* which served as a positive control (Figures S4B-F). We first demonstrated that *NEUROD1, NEUROD6, NGN1 and NGN2* were induced in the absence of MED24 and MED25 (Figures S4B-S4E). Following induction of TF expression for 2 weeks, we found that while MED24 and MED25 were dispensable for NEUROD6 mediated neuronal differentiation (Figure 6E) these factors were required for NEUROD1 and NGN1-NGN2 mediated neuronal differentiation (Figures 6F and 6G). These results show that MED24 and MED25 are required by specific proneural factors to coordinate the expression of their target genes.

**Figure 6.**
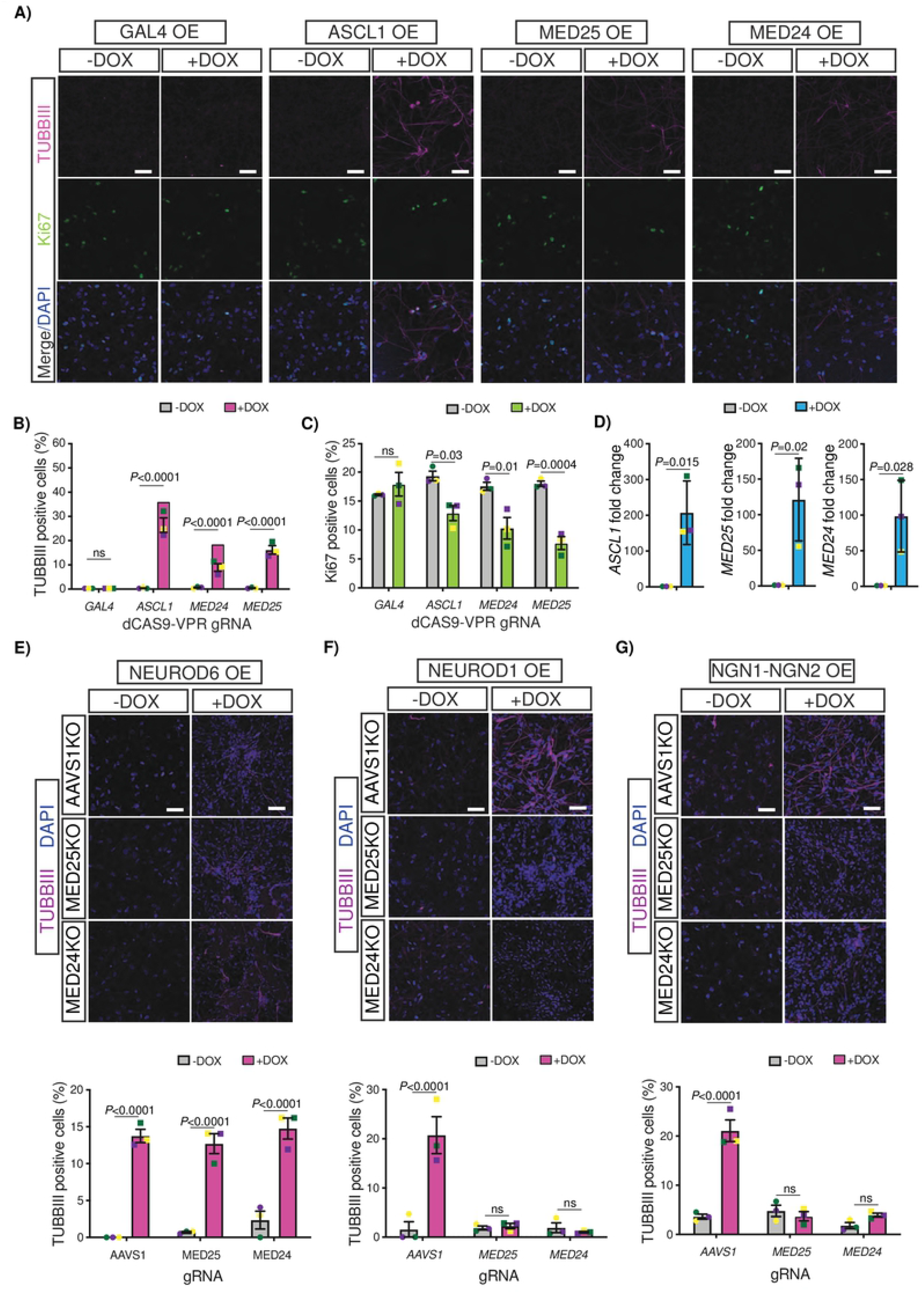
MED24-MED25 is required for bHLH transcription factors mediated neuronal reprogramming. (A) Immunocytochemical staining of TUBBIII and Ki67 upon induction of *GAL4, ASCL1, MED24* and *MED25* in GSC-dCAS9-VPR cells with DOX for 2 weeks. Cell nuclei were stained with DAPI. Scale bar, 50 μm. (B) Quantification of TUBBIII positive GSC-dCAS9-VPR cells transduced with gRNA leading to induced expression of *GAL4, ASCL1, MED24* and *MED25* following DOX treatment for 2 weeks (n=3, color-coded). two-way ANOVA and Tukey’s multiple comparison tests. Error bars, mean ± SEM. (C) Quantification of Ki67 positive GSC-dCAS9-VPR cells transduced with gRNA leading to induced expression of *GAL4, ASCL1, MED24* and *MED25* following DOX treatment for 2 weeks (n=3, color-coded). two-way ANOVA and Tukey’s multiple comparison tests. Error bars, mean ± SEM. (D) qPCR analysis of *ASCL1, MED24, MED25* mRNA levels in GSC-dCAS9-VPR cells expressing the respective gRNA and upon induction with DOX for 2 weeks (n=3, color-coded). Two-tailed unpaired t test. Error bars, mean ± SD. (E) Immunocytochemical staining (top) and quantification (bottom) of TUBBIII and Ki67 positive cells of AAVS1KO, MED24KO and MED25KO GSC-dCAS9-VPR cells transduced with NEUROD6 gRNA and induced with DOX for 2 weeks. Cell nuclei were stained with DAPI. Scale bar, 50 μm. Shown are mean values ± SEM (n=3, color-coded) tested by two-way ANOVA and Tukey’s multiple comparison tests. (F) Immunocytochemical staining (top) and quantification (bottom) of TUBBIII and Ki67 positive cells of AAVS1KO, MED24KO and MED25KO GSC-dCAS9-VPR cells transduced with NEUROD1 gRNA and induced with DOX for 2 weeks. Cell nuclei were stained by DAPI. Scale bar, 50 μm. Shown are mean values ± SEM (n=3, color-coded) tested by two-way ANOVA and Tukey’s multiple comparison tests. (G) Immunocytochemical staining (top) and quantification (bottom) of TUBBIII and Ki67 positive cells of AAVS1KO, MED24KO and MED25KO GSC-dCAS9-VPR GSCs transduced with NGN1-NGN2 gRNA and induced with DOX for 2 weeks. Cell nuclei were stained by DAPI. Scale bar, 50 μm. Shown are mean values ± SEM (n=3, color-coded) tested by two-way ANOVA and Tukey’s multiple comparison tests. See also Figure S4.

Overall, our results indicate that MED24 and MED25 play an important role in neuronal fate specification downstream of ASCL1 and other neurogenic factors in both developmental and disease contexts. Moreover, our data suggests that these Mediator subunits are sufficient for induction of neuronal differentiation revealing an underappreciated role of the Mediator tail module in this process.

## Discussion

The Mediator is a highly conserved and ubiquitously expressed protein complex in eukaryotes functioning as a core component of the RNA polymerase II transcriptional machinery. Mediator tail subunits bind to activators and repressors to modulate the context dependent transcription of genes by Pol II in various tissues (Asturias et al., 1999; El Khattabi et al., 2019; Hamilton et al., 2019; Quevedo et al., 2019; Yin and Wang, 2014). Consistent with this, an examination of our previously performed genome-wide CRISPR-Cas9 gene fitness screens in patient-derived GSC and human fetal neural stem cell lines (HFNS) (MacLeod et al., 2019; Richards et al., 2021) revealed that while head and middle subunits are essential fitness genes across all cell lines, the tail subunits including *MED24, MED25* and *MED16* show variable essentiality supporting the notion that these subunits fullfil context-specific roles in governing cellular proliferation perhaps, by controlling context-dependent transcriptional programs directing processes such as self renewal and cell differentiation (Figure S1E).

In the present study, we revealed that two Mediator complex subunits belonging to the tail module, MED24 and MED25, play essential roles in regulating cell proliferation and differentiation, governing neuronal fate identity. Upon CRISPR-Cas9 mediated knockout of either MED24 or MED25 neuronal differentiation driven by overexpression of neurogenic transcription factors or withdrawal of growth factors is abrogated. We go on to show that MED24 and MED25 are both necessary and sufficient for this process. Furthermore, using RNA-seq and ChIPseq, we demonstrated that MED24/25 cooperate with ASCL1 and other proneural transcription factors through interactions with enhancers associated with neuronal genes. ASCL1 and NGN2 bind to divergent genomic loci to induce specific neuronal fates but share in the regulation of subsets of neuronal genes(Aydin et al., 2019). Among this subset are a number of genes identified in this study to be regulated by MED25 by ChIP-Seq (*ESRRB, KLF4, TUBB3, MAP2, MYT1L*) and/or RNA-Seq (*SOX2, NES*). This is consistent with our finding that MED24/MED25 are required for both ASCL1 and NGN2 induced neuronal differentiation and supporting a major role for the Mediator tail module in neuronal fate specification. Similarly, it was recently shown that MED24 is also required for the transition from the pluripotent epiblast state to primitive endoderm identity as a result of ERK activation. This change is driven by direct modulation of enhancers associated with both pluripotency and MAPK mediated differentiation genes highlighting the importance of the tail subunits of Mediator in coordinating the specific deployment of transcriptional programs (Hamilton et al., 2019). These results suggest the possibility that the tail subunits of Mediator govern directed cell differentiation by mediating the recruitment of lineage determining transcription factors to appropriate context dependent genes thereby regulating their expression.

These results are consistent with the more recent evolutionary acquisition of Mediator tail subunits in higher eukaryotes, presumably to underlie and/or modulate specialized biological activities such as neurodevelopment (Bourbon, 2008; Conaway and Conaway, 2011). A role for the Mediator complex as a master regulator of neural stem cell identity was recently described, where Mediator was shown to colocalize with neurogenic transcription factors at enhancer and super-enhancer loci regulating their activity (Quevedo et al., 2019). Importantly, many of these TFs in the identified Mediator interaction network are known to mediate NSC self-renewal and differentiation capacity and include E-box proteins consistent with our observation of enrichment for E-box motifs at MED25 bound loci. Our study builds on this knowledge by identifying specific Mediator subunits important for neuronal differentiation and genes regulated by this complex. Systematic study of colocalization of MED24/25 and other tail subunits with these neurogenic TFs in GSCs and differentiated cells would be valuable for further understanding how this complex regulates cell identity. Many of the proteins shown to interact with Mediator in NSCs are poorly characterized, highlighting that more work is needed to fully understand the roles of Mediator and its functional relationship with interacting proteins (Quevedo et al. 2019).

While multiple studies have shown the neurogenic activity of ASCL1 overexpression, the direct examination of downstream events underlying this process has been lacking. This information is important in the context of developing directed differentiation protocols for cancer treatment (ie glioblastoma) or to control neural reprogramming for regenerative medicine applications. Another important question that remains outstanding is whether GSCs that have been differentiated towards the neuronal lineage are vulnerable to cell-cycle re-entry as is the case for astrocytic differentiation (Carén et al., 2015) or are in fact terminally differentiated as has been hypothesized (Park et al., 2017). Future studies will seek to further refine our knowledge of the latent neurogenic capacity of GSCs and identify how differentiation blockades can be overcome by modulation of neurogenic factors expression and/or activity with the goal of promoting terminal differentiation along the neuronal lineage and opposing tumorigenicity.

## Acknowledgements

The authors wish to thank Kin Chan at the Network Biology Collaborative Centre (nbcc.lunenfeld.ca) for next-generation sequencing service. Network Biology Collaborative Centre is a facility supported by Canada Foundation for Innovation, the Ontario Government, and Genome Canada and Ontario Genomics (OGI-139). We would also like to thank all members of the Angers and Dirks labs for helpful discussion throughout the course of this study. We especially thank Dr. Maira Pedroso-De-Almeida, for providing helpful feedback on the manuscript. Research was supported by the Canadian Institute for Health Research (CIHR) to SA (PJT-169054); Canadian Epigenetics, Environment, and Health Research Consortium (CEEHRC) initiative from the CIHR (TGH-158221) to SA and PBD; Stand Up to Cancer Canada Cancer Stem Cell Dream Team Research Funding (SU2C-AACR-DT-19-15) provided by the Government of Canada through Genome Canada and the CIHR, with supplemental support from the Ontario Institute for Cancer Research, through funding provided by the Government of Ontario. Stand Up To Cancer Canada is a Canadian Registered Charity (Reg. # 80550 6730 RR0001). Research Funding is administered by the American Association for Cancer Research International - Canada, the Scientific Partner of SU2C Canada.

## Author Contributions

Conceptualization M.A., G.M. and S.A.; Methodology M.A., G.M., N. R and S.A.; Formal Analysis M.A., G.M. and Z.S.; Investigation. M.A., M.K. F.M, S.L., N.I.P. and A.Y.; Resources P.B.D; Writing - Original Draft M.A., G.M. and S.A.; Writing - Review and Editing M.A., G.M. and S.A.; Supervision S.A., P.B.D.; Funding Acquisition S.A., P.B.D.

## List of Supplemental files to add

1. Supplemental file 1: CRISPR screen results-DrugZ and MAGEcK

2. Supplemental file 2: DeSeq2 results

3. Supplemental file 3: ChIPseq peaks w annotation

4. Table S1: gRNA sequences used in this study

5. Table S2: List of qPCR primers used in this study

See also Supplemental File-3.

## Supplemental figure legends

**Figure S1.**
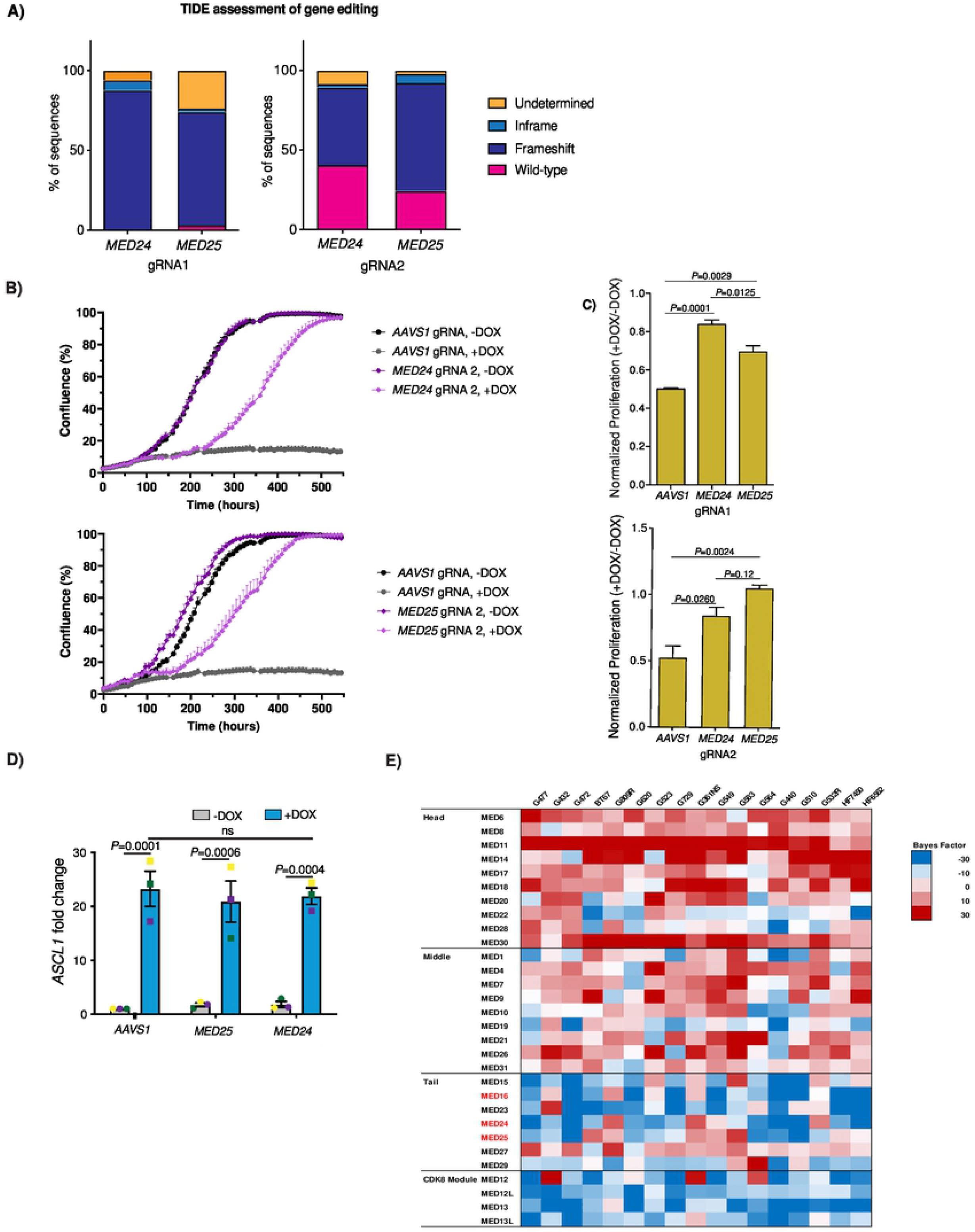
MED24 and MED25 mediate the effect of ASCL1 overexpression on induction of neuronal differentiation with a concomitant decrease in cell proliferation. Related to Figure 1. **S1A)** Quantification of CRISPR-Cas9 mediated genome-editing using gRNAs for MED24 and MED25 using the TIDE algorithm. **S1B)** Quantification of cell confluency of AAVS1KO, MED24KO (top) and MED25KO (bottom) gRNA-2 in G523-Tet-ON-ASCL1 GSCs in the presence or absence of DOX (n=3). Error bars, mean ± SD. **S1C)** Relative proliferation rate (+DOX/-DOX) of G523-Tet-ON-ASCL1 GSCs expressing the indicated gRNA-1 (top) and gRNA-2 (bottom). One-way ANOVA and Tukey’s multiple comparison tests (n=3). **S1D)** qPCR analysis of *ASCL1* mRNA levels of AAVS1KO, MED24KO and MED25KO G523-Tet-ON-ASCL1 GSCs treated with or without DOX (n=3, color-coded). Statistics from two-way ANOVA and Bonferroni’s multiple comparison tests. Error bars, mean ± SD. **S1E)** Heat map of normalized gene fitness Bayes factor (BF) scores for mediator complex from previously performed screens in patient-derived GSC lines (MacLeod et al 2019, Richards et al 2021). Subunits identified in the present study are highlighted in red.

**Figure S2.**
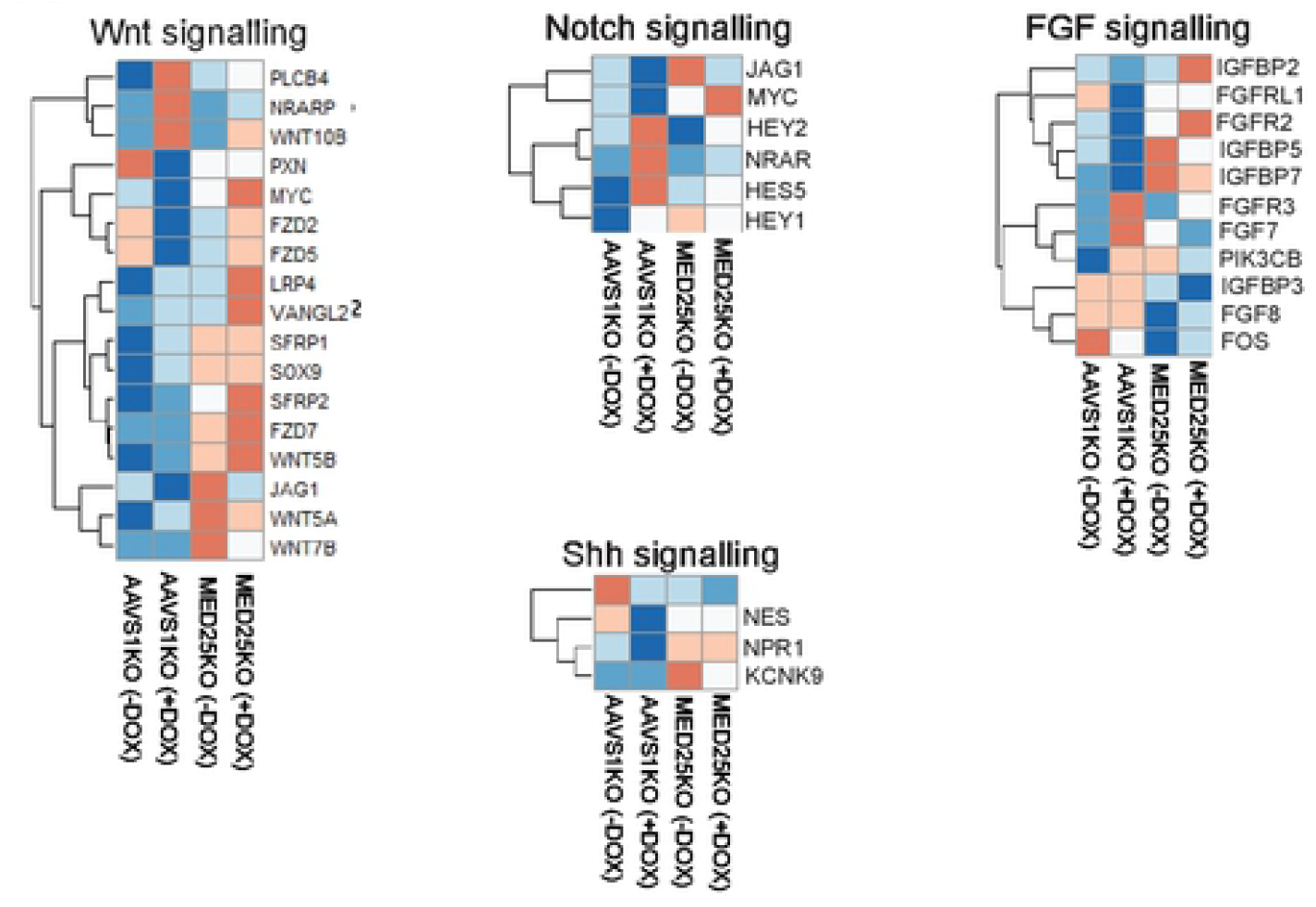
RNA-Seq analysis uncovers MED25 neuronal target genes upon ASCL1 over-expression. Related to Figure 3. **S2A)** Heatmap of transcriptional response of Wnt, Shh, Notch and FGF signaling across RNA-seq experiments. Color scale = gene-centred Z-score.

**Figure S3.**
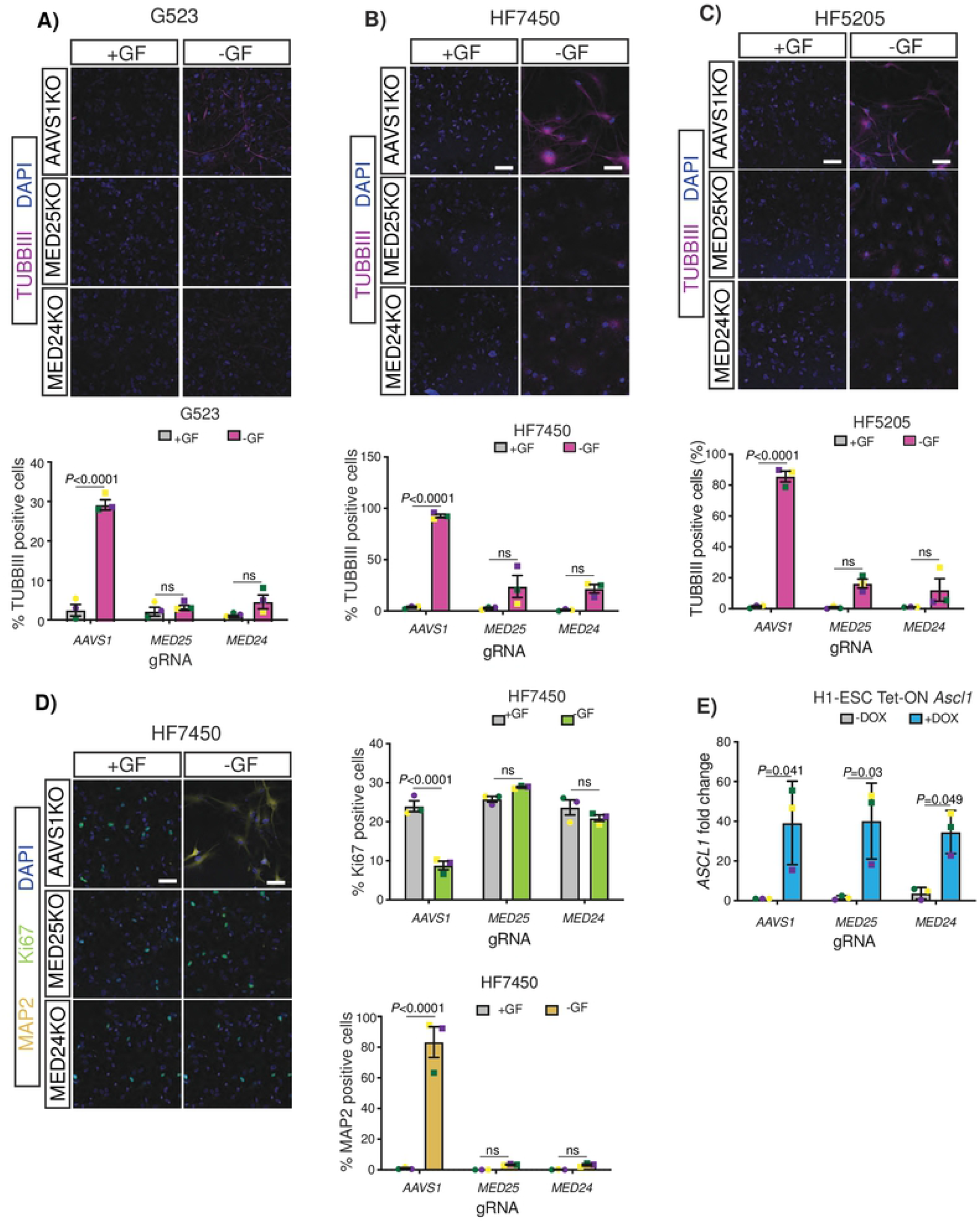
MED24 and MED25 are required for growth factor mediated neuronal differentiation in HFNS and GSCs. Related to Figure 5. **S3A)** Immunocytochemical staining (top) and quantification (bottom) of TUBBIII positive cells of AAVS1KO, MED24KO and MED25KO GSCs (G523) grown in the presence or absence of GF. Cell nuclei were stained by DAPI. Scale bar, 50 μm. Shown are mean values ± SEM (n=3, color-coded) tested by two-way ANOVA and Tukey’s multiple comparison tests. **S3B)** Immunocytochemical staining (top) and quantification (bottom) of TUBBIII positive cells of AAVS1KO, MED24KO and MED25KO HFNS (HF7450) grown in the presence or absence of GF. Cell nuclei were stained by DAPI. Scale bar, 50 μm. Shown are mean values ± SEM (n=3, color-coded) tested by two-way ANOVA and Tukey’s multiple comparison tests. **S3C)** Immunocytochemical staining (top) and quantification (bottom) of TUBBIII positive cells of AAVS1KO, MED24KO and MED25KO HFNS (HF5205) grown in the presence or absence of GF. Cell nuclei were stained by DAPI. Scale bar, 50 μm. Shown are mean values ± SEM (n=3, color-coded) tested by two-way ANOVA and Tukey’s multiple comparison tests. **S3D)** Immunocytochemical staining (left) and quantification (right) of Ki67 and MAP2 positive cells of AAVS1KO, MED24KO and MED25KO HFNS (HF7450) grown in the presence or absence of GF. Cell nuclei were stained by DAPI. Scale bar, 50 μm.Shown are mean values ± SEM (n=3, color-coded) tested by two-way ANOVA and Tukey’s multiple comparison tests. **S3E)** qPCR analysis of *ASCL1* mRNA levels of AAVS1KO, MED24KO and MED25KO ESC-Tet-ON-ASCL1 cells treated with or without DOX (n=3, color-coded). two-way ANOVA and Bonferroni’s multiple comparison tests. Error bars, mean ± SD.

**Figure S4.**
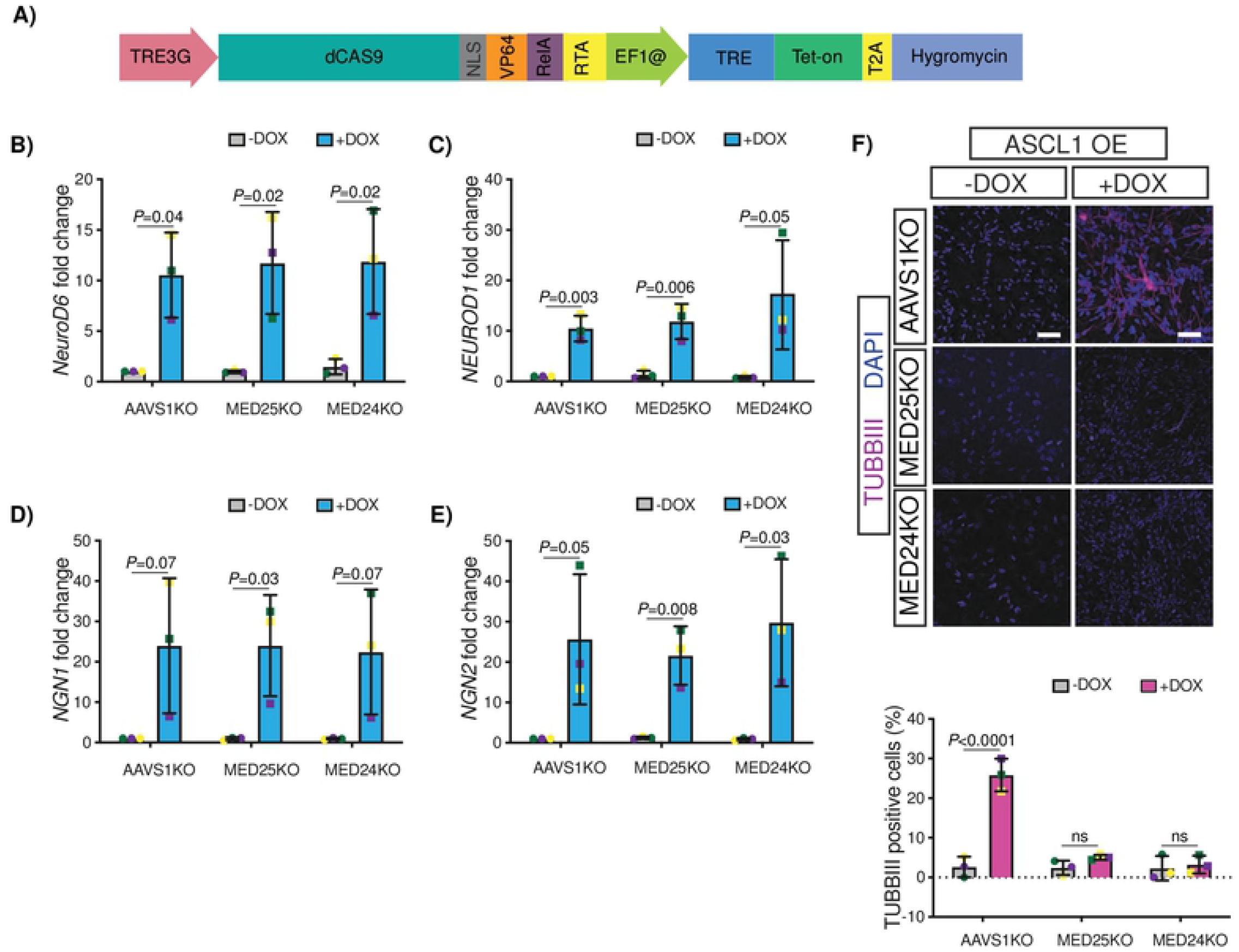
MED24 and MED25 mediate induction of neuronal differentiation upon neurogenic factor activation without impacting their expression. Related to Figure 6. **S4A)** A diagram representing dCAS9-VPR construct **S4B-E)** qPCR analysis of *NEUROD6, NEUROD1, NGN1* and *NGN2* mRNA levels upon CRISPR-activation using corresponding TSS targeted gRNAs in AAVS1KO, MED24KO and MED25KO GSC-dCAS9-VPR cells induced with DOX for 2 weeks (n=3, color-coded). Two-tailed unpaired t test. Error bars, mean ± SD. **S4F)** Top: Immunocytochemical staining of TUBBIII and Ki67 positive cells of AAVS1KO, MED24KO and MED25KO GSC-dCAS9-VPR cells infected with ASCL1 gRNA and induced with DOX for 2 weeks. Cell nuclei were stained by DAPI. Scale bar, 50 μm. Bottom: Quantification of TUBBIII positive cells (n=3, color-coded). two-way ANOVA and Tukey’s multiple comparison tests. Error bars, mean ± SEM.

## STAR Methods

### RESOURCE AVAILABILITY

#### Materials Availability

Requests for materials/reagents described in this study may be directed to, and will be fulfilled by the Lead Contact, Stephane Angers.

#### Data And Code Availability

The accession number for the RNA-seq data reported in this study is GEO:GSE166979. The accession number for the ChIP-seq data reported in this study is pending

### EXPERIMENTAL MODEL AND SUBJECT DETAILS

#### Cell culture

All samples were obtained following informed consent from patients. All experimental procedures were performed according to the Research Ethics Boards at The Hospital for Sick Children (Toronto, Canada) and the University of Toronto. Glioblastoma stem cells (G472, G523) and HFNS (HF5205, HF7450) were maintained adherently on poly L ornithine (Sigma-Aldrich) and laminin (Sigma-Aldrich) coated plates. Cell were cultured in Neurocult NS-A Basal media (StemCell Technologies) supplemented with 2 μg/mL heparin, 150 μg/mL BSA, GlutaMAX (Life Technologies), N2 supplement (Life Technologies), B27 supplement (Life Technologies), 10 ng/mL EGF (Life Technologies), 10 ng/mL FGF (Life Technologies) as described previously (Pollard et al., 2009). Cell dissociation was performed using Accutase (Life Technologies).

H1 hESCs were grown on Geltrex-coated (1:100 in DMEM/F12; Gibco) plates in StemFlex basal medium (Gibco) containing 1% Pen-Strep (Gibco). Cells were dissociated and neutralized using TrypLE Select enzyme (Gibco) and STOP solution (10% FBS in DMEM/F12) respectively. Cells were then plated onto Geltrex-coated plates in StemFlex containing RevitaCell (Gibco). RevitaCell was removed one day post seeding and the media was changed every two days. The neuronal differentiation experiments all were conducted on G-banded karyotyped H1s cultured in feeder-free and monolayer conditions.

All cell lines were maintained at 37 °C and 5% CO_2_. All cell lines were authenticated with STR profiling at The Centre for Applied Genomics (Toronto, Ontario, Canada).

#### Cell line generation

PiggyBac transposon inducible PB-Tet-ON-ASCL1 expression construct was a gift from Dr. Peter Dirks laboratory (Park et al., 2017). To enable the genome wide CRISPR screen using a gRNA library containing a puromycin resistance gene, this gene in the PB-Tet-ON-ASCL1 was replaced by Hygromycin using *FseI* and *ClaI* sites. PB-Tet-ON-ASCL1 (4 μg) was co-delivered with piggyBac transposase Hybase (1 μg) (System Biosciences) per 2 ×10^6^ patient-derived G523-GSC line using Neon nucleofection. Cells were then selected with 20 μg/mL of hygromycin. After establishment of a polyclonal stable cell line, single clones were isolated by limited dilution. These cells are henceforth referred to as G523-Tet-ON-ASCL1. Clones were expanded and tested for induction of ASCL1 upon addition of 1 μg/mL of doxycycline hyclate to NS media using qPCR and Immunofluorescence for TUBBIII as neuronal marker.

To generate stable H1 hESC PB-Tet-ON-ASCL1 expressing cell line, 1×10E^6^ cells were electroporated with 3 μg of PB-Tet-ON-ASCL1 expression vector (PB-TRE-dCas9-VPR, addgene # 63800) and 1 μg of transposase Hybase using the Neon nucleofection. Following electroporation, cells were plated onto 6-well Geltrex-coated culture dishes in the presence of RevitaCell for one day and then removed RevitaCell and allowed cells to recover for two days before selecting for PB-Tet-ON-ASCL1 expressing cells using 20 μg/mL hygromycin.

To generate stable G523-dCas9-VPR expressing cell line, 1×10E^6^ cells were electroporated with 3 μg of dCas9-VPR piggyBac expression vector (PB-TRE-dCas9-VPR, Addgene # 63800) and 1 μg of transposase Hybase (System Biosciences) using the Neon nucleofection (Chavez et al., 2015). Media was changed with fresh media the next day and 36 hours post-electroporation selection was done using 20 µg/mL hygromycin.

#### CRISPR-Cas9 gene editing

For generation of single gene knockout, individual gRNAs were ligated into *BsmBI* digested LentiCRISPRV2 (Plasmid Addgene catalogue #52961) vector. For gene activation studies, individual gRNAs selected from Human Genome-wide CRISPRa-v2 libraries (Addgene Pooled Libraries #83978, (Horlbeck et al., 2016)) were ligated into *BstXI*-*BlpI* digested hCRISPRa-v2. To transduce the cells, GSCs, HFNS as well as ESCs were infected with LentiCRISPRV2 or CRISPRa-v2 for gene knockout or gene activation, respectively by transducing 1.5×10^7^ cells in the presence of 0.8 μg/mL polybrene. Transduced cells were selected with 2 μg/mL puromycin for 2-3 days prior to seeding for performing various experiments. TIDE (Brinkman et al., 2014) was used to quantitatively assess gene editing from PCR amplicons flanking gRNA target sites. For a complete list of gRNA sequences, see Table S1.

#### Lentivirus production

Lentivirus containing TKOv3 gRNA library (Addgene Pooled Library #90294) was prepared as described (Hart et al., 2017) which was then concentrated using Lenti-X Concentrator solution (Clontech) according to the manufacturer’s protocol. For small-scale lentivirus production of individual gRNA, 4×10^6^ HEK293T cells were seeded on 10 cm plates and the next day transfected with 5 μg lentiviral plasmid, 5 μg of psPAX2 and 2 μg of pMD2.G. Media was changed 24 hours post-transfection and viral media was harvested 48 hours post-transfection. Viral supernatant was cleared by centrifugation at 1,000 x g for 5 minutes and filtered and concentrated using Lenti-X Concentrator solution.

### METHOD DETAILS

#### Genome-wide CRISPR-Cas9 screen

Genome-wide CRISPR-Cas9 knockout screen was performed using the previously described 70K TKOv3 library (Hart et al., 2017; MacLeod et al., 2019). 500×10^6^ G523-Tet-ON-ASCL1 cells were transduced at an M.O.I. of 0.3-0.4 in the presence of 0.8 μg/mL polybrene and plated onto sixty 10-cm dishes. Twenty-four hours post-transduction, media was replaced with fresh Glioblastoma Stem Cell media containing 2 μg/mL puromycin. After 3 days of puromycin selection, cells were combined and 2×10^7^ cells were collected as T0 samples and stored at -80°C for later processing. The pool of remaining cells were divided equally into 3 replicates each for DMSO or DOX to induce ASCL1 expression for 4 weeks. At all passages a minimum of 200-fold library coverage was maintained for all replicates. After 4 weeks of culture cells were collected for gDNA extraction.

#### Genomic DNA Library Preparation and Sequencing

Genomic DNA was extracted using the QiaAMP DNA Blood Maxi Kit (QIAGEN) according to the manufacturer’s protocol. For each gDNA sample, 50μg of gDNA was processed as previously described (MacLeod et al., 2019) for Illumina sequencing using unique combinations of i5 and i7 barcode sequences for each sample. Barcoded PCR product was then gel purified and sequenced using a NextSeq500 instrument (Illumina) at read depths of 500- and 200-fold for T0 and other samples respectively.

#### Tissue and Cell Staining and Microscopy

GSCs and HFNS Cells were fixed for 15 minutes with 4% PFA and either processed immediately or stored at 4°C for up to one week in PBS for later processing. Cells were permeabilized in 0.2% Triton X-100 PBS, washed twice in PBS and were blocked using 3% normal goat serum containing 0.1% Triton-X for 1 hour at room temperature. Primary antibody concentration was prepared in blocking solution and were incubated overnight at 4°C. Cells were then washed three times with PBS and incubated with Fluorophore-conjugated secondary antibodies at 1:500 dilution in blocking solution for 1 hour at room temperature followed by several washes. Coverslips were mounted using Fluoromount (Sigma-Aldrich) containing DAPI. Images were captured using 20x/0.8 objective on Zeiss LSM700 confocal microscope operated with ZEN software black edition.

ESCs were fixed for 15 minutes with 4% PFA and cell permeabilization was performed in 0.3% Triton X-100 for 10 minutes followed by blocking in 1% BSA for 1 hour at room temperature. Cells were then incubated with primary antibodies overnight at 4°C, followed by three washes with PBS and 1 hour incubation with Fluorophore-conjugated secondary antibodies at 1:500 dilution at room temperature. Cells were mounted and analyzed as described above.

Ki67 (PA5-16785, Invitrogen, 1:500 dilution), TUBBIII (TU-20, Millipore-Sigma, 1:500 dilution), Nestin (Cat#AB5922, Millipore, 1:800 dilution), MAP2 (Cat#MAB3418, Millipore, 1:500 dilution), GFAP (PA5-16291, Thermo fisher scientific, 1/300 dilution) and OCT3/4 (sc5279, Santa Cruz, dilution 1:100) primary antibodies were used in this study. Alexa Fluor 488-labeled donkey anti-rabbit (A21206, Invitrogen) or Alexa Fluor 594-labeled donkey anti-mouse Ab (A21203, Invitrogen) were used as secondary antibodies.

#### *In Vitro* GSC Differentiation Assay

GSC differentiation experiments were either performed by sequential withdrawal of rhEGF and FGF or by induction of ASCL1 using DOX treatment as previously described (Park et al., 2017; Pollard et al., 2009; Rajakulendran et al., 2019). Briefly, for GF withdrawal, cells were seeded in Glioblastoma Stem Cell media and 1-2 days post-seeding, media was changed to NS media containing reduced bFGF (5 ng/mL) and lacking rhEGF. 7-8 days post-seeding, media was changed to Glioblastoma Stem Cell media lacking both rhEGF and bFGF, and cells were maintained in culture under this condition for 2 more weeks. ASCL1-mediated neuronal differentiation experiments were performed by culturing cells in NS media for 1 day followed by changing media to NS media containing either DMSO as control or 1 µg/mL of DOX for 14 days. Media was changed every 2 days.

#### *In Vitro* ESCs Differentiation Assay

The H1 hESC PB-Tet-ON-ASCL1 expressing cell line was seeded in 24-well Geltrex-coated plates in the presence of RevitaCell for one day which was then replaced by fresh StemFlex media containing either DMSO as control or 0.8 µg/mL DOX. The media was changed every two days for 7 days.

#### *In Vitro* Cell Proliferation Assay

Cells were seeded adherently on a 96-well plate in biological triplicate in Glioblastoma Stem cell media for 1 day. Next day, cells were treated with either DMSO as control or 1 µg/mL DOX to induce ASCL1 expression, for three weeks. *CellTiter*-*Glo*^®^ Luminescent Cell Viability Assay (Promega) was then used according to the manufacturer’s protocol to assess the cell viability.

#### *In Vitro* Live-Cell Imaging

Cells were plated on coated dishes in biological triplicates on a 96-well plate and imaged using IncuCyte S3. Cells were imaged using phase-contrast every 16 hour over a 22 day period. Cell confluency measured using IncuCyte live cell analysis system.

#### *In Vitro* Limiting Dilution Assay

*In vitro* sphere-forming ability was measured by culturing cells in serial dilutions (range of 3–2000) on non-adherent 96-well plates under NS conditions. After 10 days of culture, the frequency of sphere-forming cells was determined by inequality in frequency between multiple groups and tested for adequacy of the single-hit model using Extreme limiting dilution analysis (ELDA) software (http://bioinf.wehi.edu.au/software/elda) (Hu and Smyth, 2009).

#### RNA Sequencing

Cells were grown on coated plates for 1 week. Total RNA was isolated from cells using the TRIzol reagent (Thermofisher). The quality of RNA was measured using Bioanalyzer (Agilent) and RNA-seq libraries (3 biological replicates each samples) were derived using TruSeq V2mRNA-enriched library kit (Illumina) according to the manufacturer’s instruction. Libraries were sequenced on an Illumina NextSeq-500 instrument with 75-bp single-end reads. The trimmed reads were aligned to the reference genome (UCSC hg19) using Kallisto (Bray et al., 2016) and differential expression analysis was performed using the DESeq2 package (Love et al., 2014). Pathway and gene ontology analyses were performed using gProfiler (Reimand et al., 2016).

#### Quantitative Polymerase Chain Reaction (qPCR)

Cells were collected and stored at -80°C. Total RNA was extracted from cells using the TRIzol reagent (Thermofisher) and cDNA synthesis was performed using 2 µg of total RNA which was reverse-transcribed using the Super-Script II reverse transcription kit (Thermo Fisher) and qPCR was performed using SyBrGreen (ThermoFisher) on BioRAD CFX instrument. The primer sequences can be found in Table S2.

#### ChIP Sequencing Sample Preperation

Adherently cultured cells were treated with DMSO as control or 1 μg/mL DOX for 18 hours in biological triplicates (15×10^6^ cells per replicate). Cells were fixed using 1% formaldehyde for 10 minutes at room temperature and neutralized by 0.65 M glycine solution, washed 3 times with 10 mL ice-cold PBS (centrifuged at 2000 rpm for 5 minutes in 4°C centrifuge) and flash frozen with liquid nitrogen after addition of protease inhibitor cocktail and stored at -80°C. Frozen fixed cell pellets were resuspended in swelling buffer (25 mM Hepes, pH 7.8, 1.5 mM MgCl2, 10 mM KCl, 0.1% NP-40, 1mM DTT, 1X protease inhibitor cocktail) and incubated on ice for 10 minutes followed by 40 times dounce homogenization. After centrifugation pellets were resuspended in sonication buffer (50mM Hepes, pH 7.9, 140 mM NaCl, 1mM EDTA, 1% Triton X-100, 0.1 % Na-deoxycholate, 0.1% SDS, 1X protease inhibitor cocktail) and nuclei were sonicated using a Biorupter sonicator with the following settings: Intensity: high; Cycles: 30 seconds on/off, Time: 30 min. The sonicated solution was transferred into low binding tubes and centrifuged at 14000 rpm for 15 minutes and the supernatant was then collected. 50 μL of each sample was retained as input and the 40 μL of Dynabeads Protein-A and G (Life Technologies) prebound to MED25 antibody (PA5-43617, Invitrogen) or IgG (Cell Signaling Technology) added to the remaining chromatin and rotated at 4°C overnight. Crude chromatin was then pre-washed with 1 mL sonication buffer followed by three times washing with 1 mL wash buffer A (50 mM Hepes pH 7.9, 500 mM NaCl, 1 mM EDTA, 1% Triton X-100, 0.1% Na-deoxycholate, 0.1% SDS, protease inhibitor) and three times washing with 1 mL wash buffer B (20mM Tris, pH 8.0, 1mM EDTA, 250mM LiCl, 0.5% NP-40, 0.5% Na-deoxycholate, protease inhibitor) at 4°C. To remove detergents and salts from the previous washes, the chromatin was washed two more times with 1 mLμ TE buffer containing 50mM NaCl. Beads were eluted and incubated at 65°C for 20 minutes, centrifuged at 14000 rpm for 1 minute and supernatant was collected by using a magnet rack. Immunoprecipitated chromatin as well as input were reverse cross-linked at 65°C for 5 hours. DNA cleanup was performed using Qiaquick PCR purification Kit (Qiagen) following manufacturer’s instructions.

Library preparation and sequencing of ChIP-seq experiments were performed by the Lunenfeld-Tanenbaum Sequencing Facility. ChIP-seq sequences libraries were prepared from 5 ng fragmented DNA using the llumina TruSeq ChIP Library Preparation Kit (Cat# IP-202-2012) according to the manufacturer’s instructions. Quality checked libraries were then loaded onto an Illumina NextSeq 500 run with Illumina NextSeq 500/550 Hi Output Kit v2.5 (75 Cycles).

### QUANTIFICATION AND STATISTICAL ANALYSIS

#### Analysis of Genome-wide CRISPR-Cas9 screen

FASTQ files were demultiplexed and trimmed of adaptor sequences prior to mapping of sequencing reads to TKOv3 library using the MAGeCK (Li et al., 2014) *test* function. Gene knockout positively selected under DOX treatment were identified using the DrugZ (Colic et al., 2019) software and MAGeCK (Li et al., 2014) *count* function, both using unpaired experimental design.

#### Analysis of ChIP-seq Data

Real-time base call (.bcl) files were converted to FASTQ files using Illumina bcl2fastq2 conversion software v2.17. The reads were aligned to the hg19 genome using bowtie2 (Langmead and Salzberg, 2012). Alignments were filtering to retain only uniquely mapped reads and remove instances of low-quality mapping. Replicates were merged and MED25-bound regions were detected using MACS2 v2.0.10 with default settings and a q-value filter of 0.05. Analysis of MED25-bound regions was performed using the Homer software package (v4.11.1) (Heinz et al., 2010) including the *annotatePeaks* package for annotation of peak proximal genes and classification of genomic regions, and the mergePeaks package for identification of overlapping and differential peaks. MED25-bound peaks were annotated as promoter peaks if they overlapped with a 3kb window (2.5kb upstream to 0.5 kb downstream) around the TSS. Cell type specific enhancer lists were obtained from Enhancer Atlas version 2.0 (Gao and Qian, 2020) and HACER (Gao and Qian, 2020; Wang et al., 2019a)

## Statistical Analysis

Statistical details related to each experiment can be found in the figure legends. Unless otherwise indicated Graphpad Prism 5 was used for the statistical analyses (GraphPad Software, La Jolla, California, USA). Statistical analysis of qPCR data was performed using GraphPad Prism and two-way ANOVA test was used to calculate significance. Values were obtained from at least 2 biological replicates for each sample and GAPDH was used for normalization. Statistical analysis of immunocytochemical data was performed in GraphPad Prism and two-way ANOVA test was used to calculate significance. Error bars represent mean ± SEM. Data was obtained from 3 biological replicates and 3 technical replicates. Live-cell imaging analysis was plotted using GraphPad Prism and error bars represent mean ± SD (n = 3 biological replicates and technical replicates). In vitro LDA was performed in ELDA software (see above) and SFC is plotted as 95% confidence intervals for 1/stem cell frequency. To assess the differences in stem cell frequencies between various groups Pairwise chi-square tests were used. Statistical analysis of proliferation data was performed in GraphPad Prism and Two-way ANOVA was used to calculate the significance. Error bars represent mean ± SEM (n = 3 biological replicates). In all figures: ^∗^, p < 0.05; ^∗∗^, p < 0.01; ^∗∗∗^, p < 0.001; ^∗∗∗∗^, p < 0.0001.

